## Supplemental Figures for "Mimicked synthetic ribosomal protein complex for benchmarking crosslinking mass spectrometry workflows"

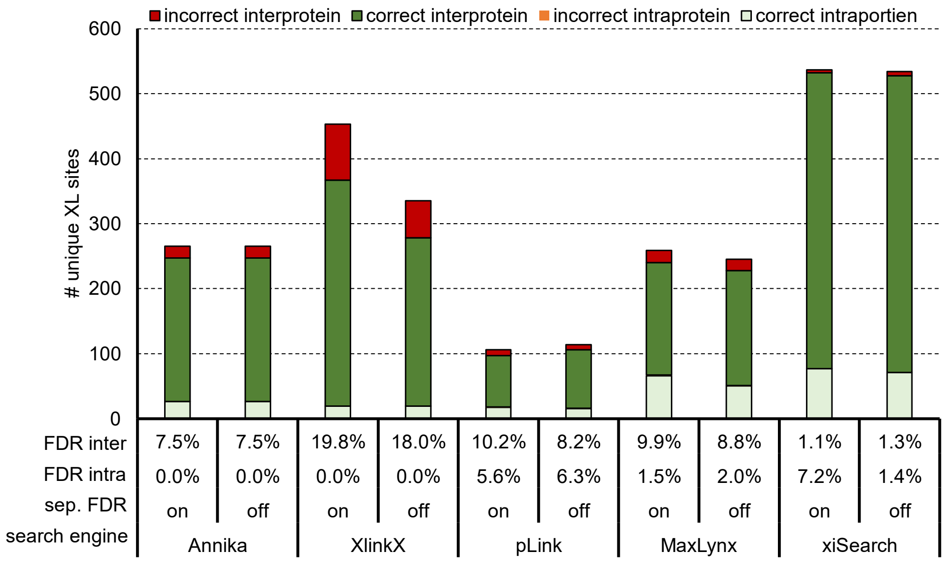


*Figure S 1:* ***Effect of separate inter/intra FDR calculation in a complex environment****. Average number of unique XL sites from the DSSO crosslinked main library spiked into non – crosslinked tryptic HEK peptides (1:5) after acquisition using a stepped HCD MS2 strategy with separate FDR calculation for inter- and intra-crosslinked peptides set on or off. A proteome wide search was performed with the indicated software. Although synthetic peptides were used for crosslinking their sequences are based on ribosomal protein sequences. "Intraprotein" links correspond to homomeric links and "interprotein" are heteromeric links based on the proteins the synthetic peptides would belong to. Experimentally validated FDR within each crosslink group is shown below, n=1.*

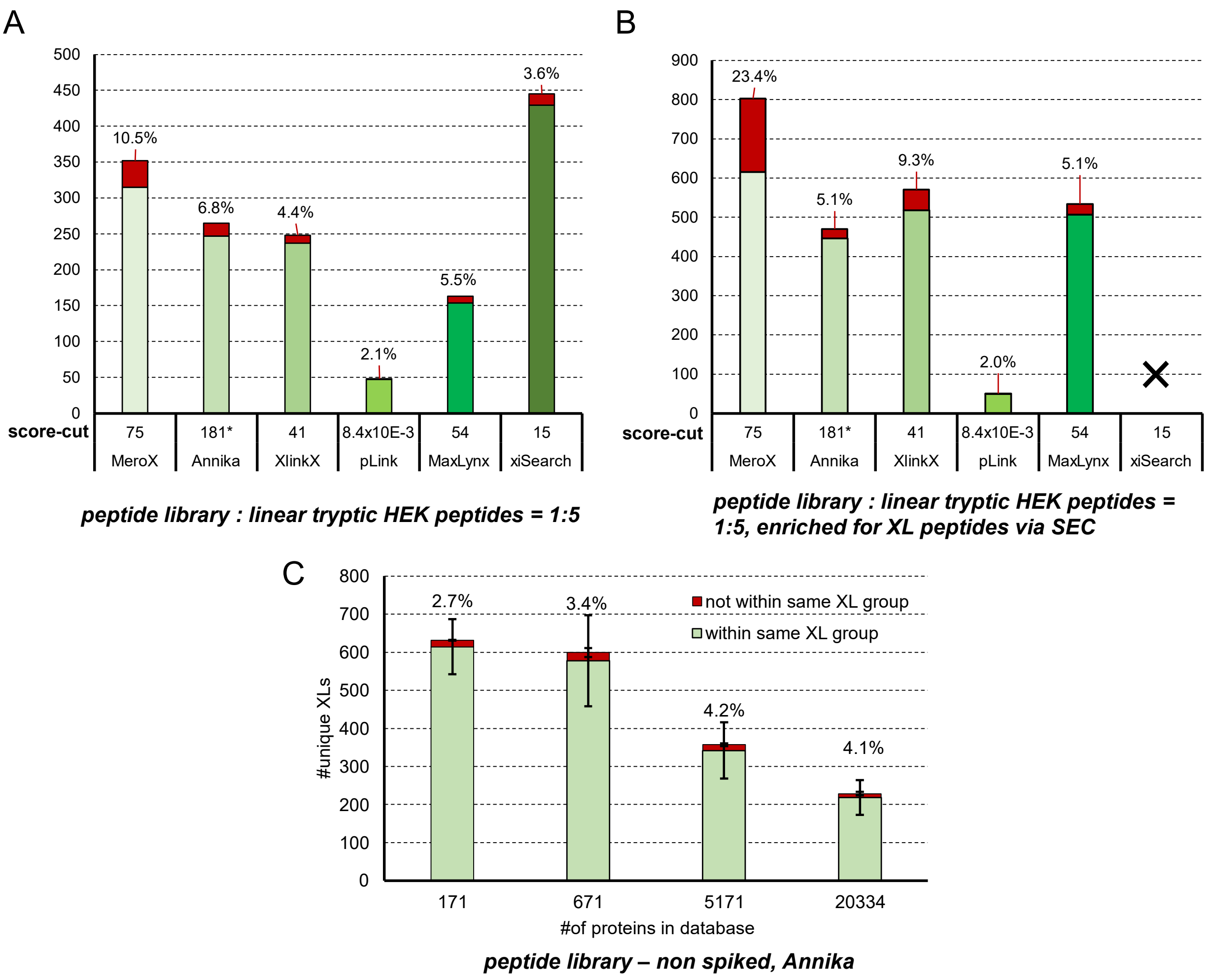


*Figure S 2:* ***Performance benchmarking in a mimicked complex environment searching against the full proteome and effect of variation of database size independent of sample complexity****. The DSSO linked main library was mixed with linear tryptic HEK peptides (1:5 w/w). Bars indicate the number of unique XL sites identified using the indicated algorithm at 1% estimated FDR and by applying the given score-cutoff values that were chosen based on experiments without spiking to reach a real FDR of 1%. The database contains 171 E. coli ribosomal proteins plus an additional 20163 human proteins. (****A****) direct measurement (****B****) measurement after enrichment for crosslinked peptides by size exclusion chromatography. Of note, analysis of the 5 SEC fractions did reproducibly not work with our largest 20334 protein database and xiSearch, as the software crashes. This data is therefore missing.* ********: 180 is the minimal score-cutoff based on our results shown in Figure 1, but the actual lowest scored link that passed the FDR threshold was 214 (A) and 256 (B) and as shown in Figure 5. (****C****) The non-spiked DSSO linked main library data as shown in figure 1A was analyzed against databases of increasing size as done in figure 5, n=3, error bars indicate standard deviations.*

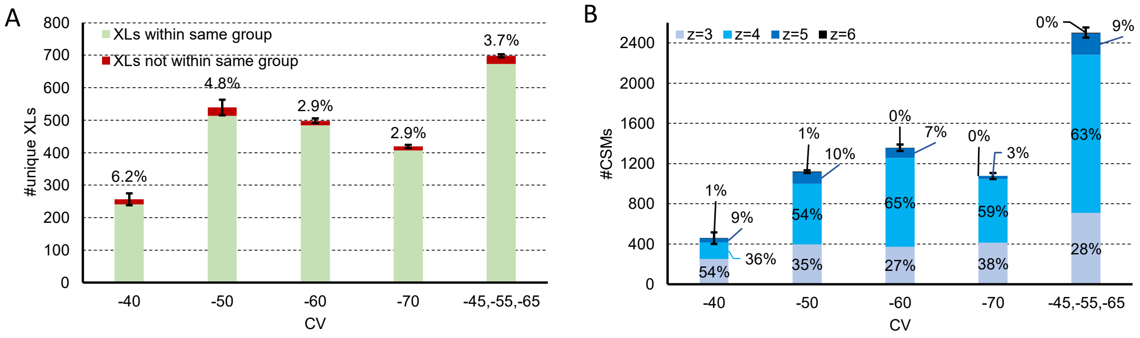


*Figure S 3:* ***DSSO linked peptides on a FAIMS equipped device.*** *800 ng of DSSO cross-linked synthetic peptides were measured under variation of the compensation voltage (CV) or using a method encompassing 3 CVs. Bars indicate unique XLs and*

*numbers above indicate the experimentally validated FDR (**A**), in panel (**B**) bars indicate the number of CSM IDs at 1% FDR and their charge distribution, error bars indicate standard deviations of average values, n=3*

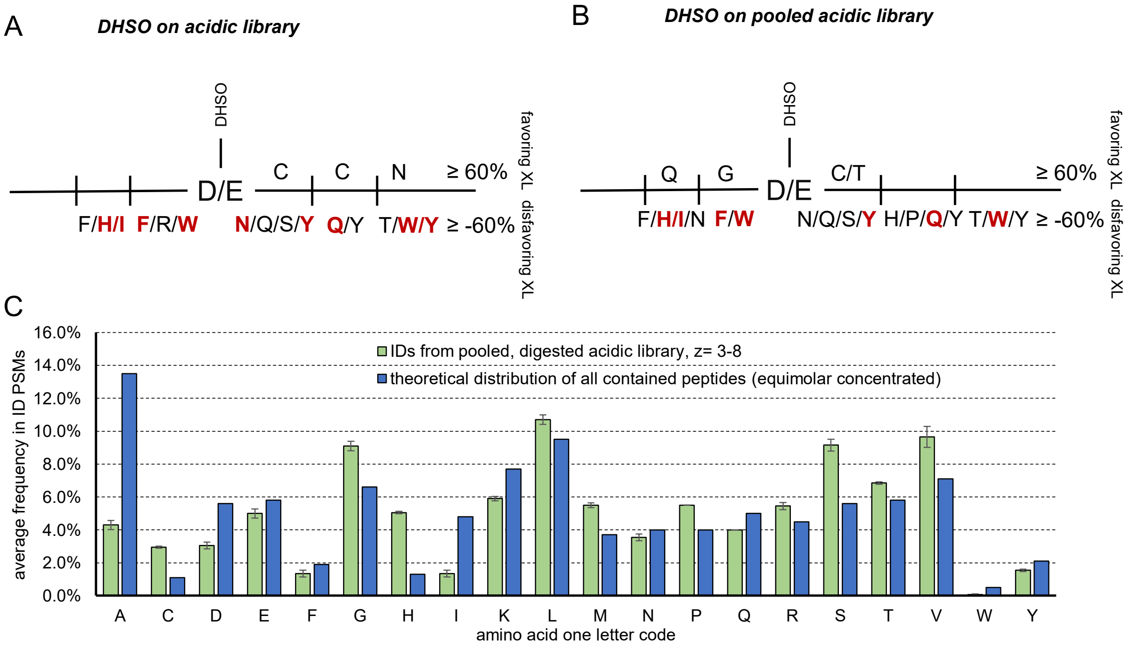


*Figure S 4: **Influence of specific amino acids in proximity to the XL-site influencing the formation of a crosslink**. Based on result from the acidic peptide library XLed with DHSO. Given % correspond to the average frequency of a specific amino acid at the given position relative to the cross-linked site within all identified CSMs. Normalized to expected frequencies within all (in silico generated) cross links of the library system. Amino-acids marked in red were not identified in any crosslink although it the corresponding peptides are existing in the peptide mix (**A**) DHSO cross-linked (in separate groups) peptides of acidic library, n=3 (B) DHSO cross-linked pool of all peptides from acidic library, n=3 (C) Amino-acid frequency in identified non-crosslinked peptides vs expected distribution.*

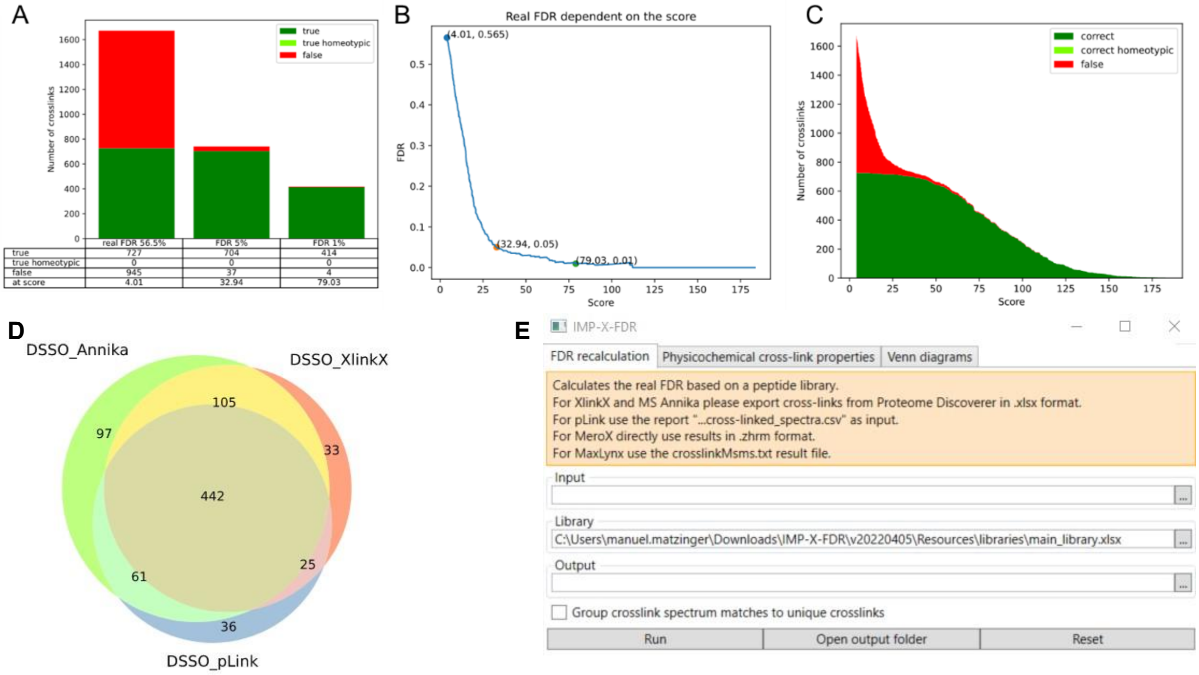


*Figure S 5: **Exemplary output figures of IMP-X-FDR**. Using the FDR recalculation functionality outputs three figures showing the number of cross-links and FDR as well as post-score cutoff corrected numbers (**A**), the real FDR vs score (**B**), and the number of crosslinks identified vs score (**C**). Using the Venn diagrams functionality, the overlap of 2 -4 datasets can be visualized (**D**). (**E**): Design of the graphical user interface.*

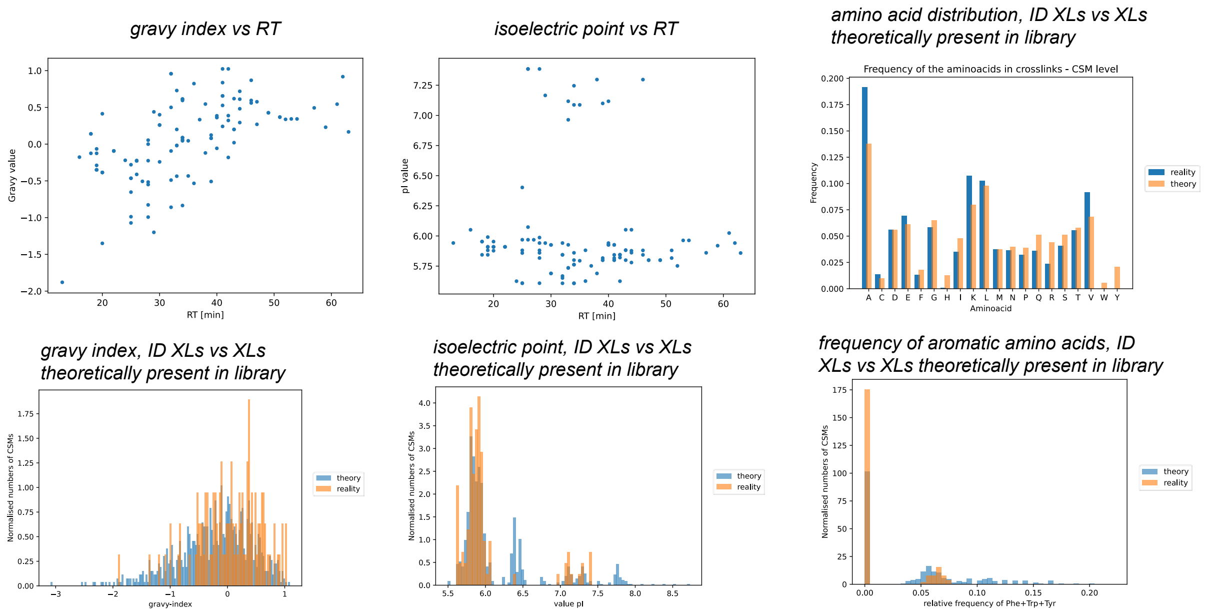


*Figure S 6:* ***Exemplary output figures of the physicochemical cross-link properties functionality of IMP-X-FDR****. IMP-X-FDR outputs graphs based on the amino acid distribution, gravy index, isoelectric point and frequency of aromatic amino acids in an automated manner.*

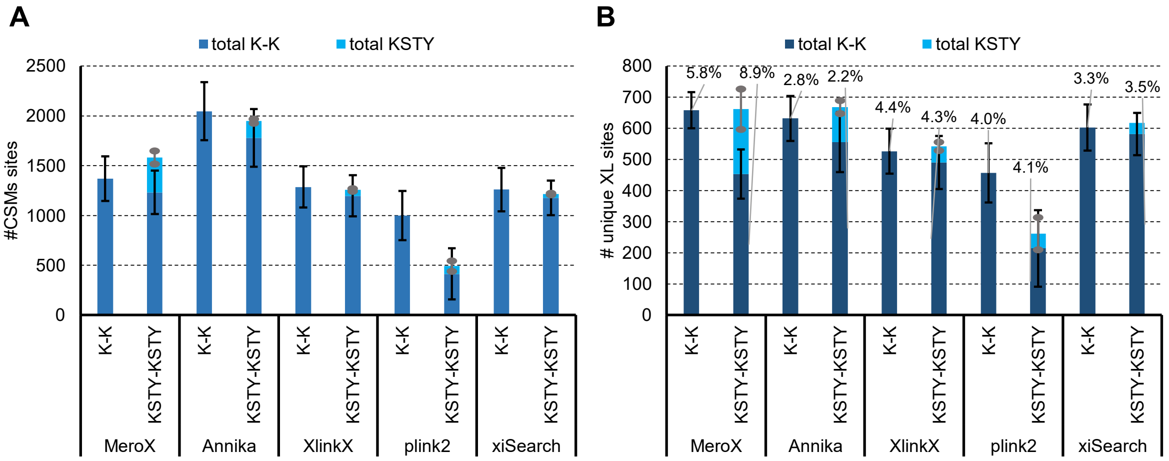


*Figure S 7:* ***Effect of site localization searches (KSTY) for FDR estimation.*** *Based on the results of the main peptide library crosslinked with DSSO. (A) Comparison of crosslink searches using only K to K (dark blue) crosslinks sites and KSTY to KSTY (light blue) on CSM level at 1% link level FDR. (B) Comparison of crosslink searches using only K to K (dark blue) crosslinks sites and KSTY to KSTY (light-blue) on unique link level at 1 % FDR. Black and grey error bars indicate standard deviations of average values for K-K and KSTY-KSTY respectively, n=3. Percentage above the bars for unique residue pairs, represent "experimentally estimated FDR".*

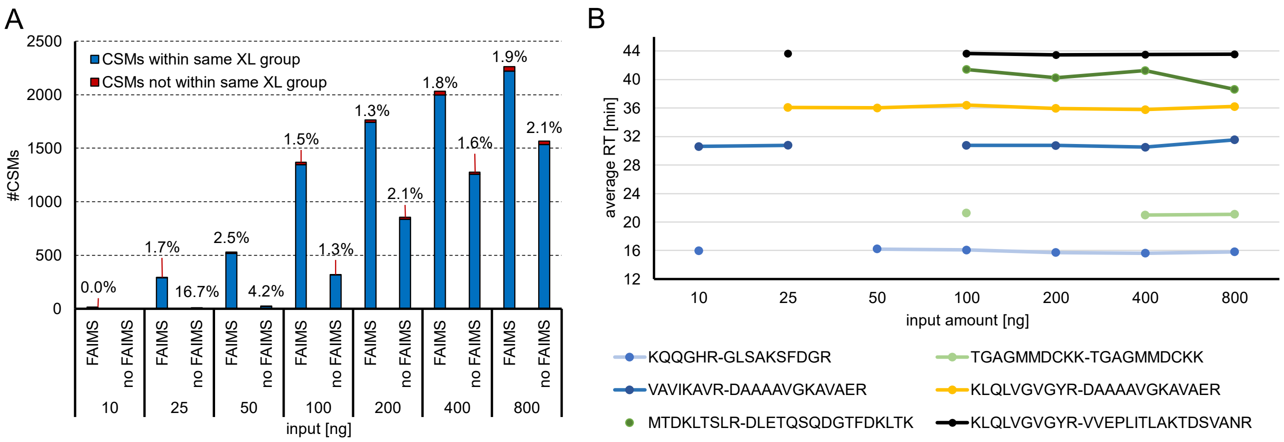


*Figure S 8:* ***Impact of FAIMS and input amount to identified CSMs****. Input amounts, as indicated, of the DSSO cross-linked main library were measured on an Orbitrap Exploris 480 using a stepped HCD MS2 strategy with or without a FAIMS device attached. Data was analyzed using Annika at 1% FDR. (**A**)Bars indicate the number of CSM matches. (**B**) Representative CSM sequences acquired with FAIMS and their retention time in dependence on input amount.*
