## Supplemental Tables for "Mimicked synthetic ribosomal protein complex for benchmarking crosslinking mass spectrometry workflows"

Supplementary Table 1: List of synthesized peptides and allocation to their crosslink group

| ID number | tryptic sequence (miscleavage at XL site) | synthesized sequence (Ac = acetyl, Am = amide, N3 = azide) | group annotation in main library | group annotation in enrichable library | group annotation in acidic library | annotated protein accession number |
|---|---|---|---|---|---|---|
| 1 | VALVAKIGENINIR | Ac-WGGGGRVALVAKIGENINIR | 1;11 | 1;9 | no | P0A6P1 |
| 2 | EIAEKMVEGR | Ac-WGGGGREIAEKMVEGR | 1;12 | 1;10 | no | P0A6P1 |
| 3 | KAGNVAADGVIK | Ac-WGGGGRKAGNVAADGVIK(N3) | 1;13 | no | no | P0A6P1 |
| 4 | EFIAKLQANPAK | Ac-WGGGGREFIAKLQANPAK(N3) | 1;14 | no | no | P27302 |
| 5 | MAALMKQR | Ac-WGGGGRMAALMKQR | 1;15 | 1;11 | no | P27302 |
| 6 | ILKCGFR | Ac-WGGGGRILKCGFR | 1;16 | 1;12 | no | P0AG44 |
| 7 | VKDLPGVR | Ac-WGGGGRVKDLPGVR | 1;17 | 1;13 | no | P0A7S3 |
| 8 | MAHIEKQAGELQEK | Ac-WGGGGRMAHIEKQAGELQEK(N3) | 1;18 | no | no | P0A7W1 |
| 9 | QAGELQEKLIAVNR | Ac-WGGGGRQAGELQEKLIAVNR | 1;19 | 1;14 | no | P0A7W1 |
| 10 | EVPAAIQKAMEK | Ac-WGGGGREVPAAIQKAMEK(N3) | 1;20 | no | no | P0A7W1 |
| 11 | VKGGFTVELNGIR | Ac-WGGGGRVKGGFTVELNGIR | 2;11 | 1;15 | no | P0AG67 |
| 12 | ITDVEVLKAQFEEER | Ac-WGGGGRITDVEVLKAQFEEER | 2;12 | 1;16 | no | P0A6P1 |
| 13 | TGAGMMDCKK | Ac-WGGGGRTGAGMMDCKK(N3) | 2;13 | no | no | P0A6P1 |
| 14 | KAGFVTR | Ac-WGGGGRKAGFVTR | 2;14 | 2;9 | no | P0A7X3 |
| 15 | HKATLLGLGLR | Ac-WGGGGRHKATLLGLGLR | 2;15 | 2;10 | no | P0AG51 |
| 16 | MAKTIK | Ac-WGGGGRMAKTIK(N3) | 2;16 | no | no | P0AG51 |
| 17 | VKHPSEIVNVGDEITVK | Ac-WGGGGRVKHPSEIVNVGDEITVK(N3) | 2;17 | no | no | P0AG67 |
| 18 | ITLNMGVGEAIADKK | Ac-WGGGGRITLNMGVGEAIADKK(N3) | 2;18 | no | no | P62399 |
| 19 | QCKANPWQQFAETHNK | Ac-WGGGGRQCKANPWQQFAETHNK(N3) | 2;19 | no | no | P0AG67 |
| 20 | ANPWQQFAETHNKGDR | Ac-WGGGGRANPWQQFAETHNKGDR | 2;20 | 2;11 | no | P0AG67 |
| 21 | AMEKAR | Ac-WGGGGRAMEKAR | 3;11 | 2;12 | no | P0A7W1 |
| 22 | IGVPFVDGGVIKAEVVAHGR | Ac-WGGGGRIGVPFVDGGVIKAEVVAHGR | 3;12 | 2;13 | no | P0AG48 |
| 23 | DTLHLEGKELEFK | Ac-WGGGGRDTLHLEGKELEFK(N3) | 3;13 | no | no | P0AG67 |
| 24 | TDKFIVR | Ac-WGGGGRTDKFIVR | 3;14 | 2;14 | no | P60422 |
| 25 | TVGQLLKEHNAEVTGFIR | Ac-WGGGGRTVGQLLKEHNAEVTGFIR | 3;15 | 2;15 | no | P0A6P1 |
| 26 | MAEITASLVKELR | Ac-WGGGGRMAEITASLVKELR | 3;16 | 2;16 | no | P0A6P1 |
| 27 | SGAIKAAK | Ac-WGGGGRSGAIKAAK(N3) | 3;17 | no | no | P0A6P1 |
| 28 | VVSMPSTDAFDKQDAAYR | Ac-WGGGGRVVSMPSTDAFDKQDAAYR | 3;18 | 3;9 | no | P27302 |
| 29 | AVEEARAVTDKPSLLMCK | Ac-WGGGGRAVEEARAVTDKPSLLMCK(N3) | 3;19 | no | no | P27302 |
| 30 | FAAYAKAYPQEAAEFTR | Ac-WGGGGRFAAYAKAYPQEAAEFTR | 3;20 | 3;10 | no | P27302 |
| 31 | DFLKHNPQNPSWADR | Ac-WGGGGRDFLKHNPQNPSWADR | 4;11 | 3;11 | no | P27302 |
| 32 | EAGQAKESAWNEK | Ac-WGGGGREAGQAKESAWNEK(N3) | 4;12 | no | no | P27302 |
| 33 | MKGEMPSDFDAK | Ac-WGGGGRMKGEMPSDFDAK(N3) | 4;13 | no | no | P27302 |
| 34 | DIDGHDAASIKR | Ac-WGGGGRDIDGHDAASIKR | 4;14 | 3;12 | no | P27302 |
| 35 | IGVLVAAKGADEELVK | Ac-WGGGGRIGVLVAAKGADEELVK(N3) | 4;15 | no | no | P0A6P1 |
| 36 | LQANPAKIASR | Ac-WGGGGRLQANPAKIASR | 4;16 | 3;13 | no | P27302 |
| 37 | VVEPLITLAKTDSVANR | Ac-WGGGGRVVEPLITLAKTDSVANR | 4;17 | 3;14 | no | P0AG44 |
| 38 | HEIIKTTLPK | Ac-WGGGGRHEIIKTTLPK(N3) | 4;18 | no | no | P0AG44 |
| 39 | TTLPKAK | Ac-WGGGGRTTLPKAK(N3) | 4;19 | no | no | P0AG44 |
| 40 | MAVQQNKPTRSK | Ac-WGGGGRMAVQQNKPTRSK(N3) | 4;20 | no | no | P0A7N4 |
| 41 | TSGEKHLR | Ac-WGGGGRTSGEKHLR | 5;11 | 3;15 | no | P0A7N4 |
| 42 | VFQTHSPVVDSISVKR | Ac-WGGGGRVFQTHSPVVDSISVKR | 5;12 | 3;16 | 1;9 | P0A7K6 |
| 43 | KISNGEGVER | Ac-WGGGGRKISNGEGVER | 5;13 | 4;9 | no | P0A7K6 |
| 44 | AKLYYLR | Ac-WGGGGRAKLYYLR | 5;14 | 4;10 | no | P0A7K6 |
| 45 | LDLYITVKGGGISGQAGAIR | Ac-WGGGGRLDLYITVKGGGISGQAGAIR | 5;15 | 4;11 | no | P0A7X3 |
| 46 | KSSAAR | Ac-WGGGGRKSSAAR | 5;16 | 4;12 | no | P0A7X3 |
| 47 | AFNAKTDSIEK | Ac-WGGGGRAFNAKTDSIEK(N3) | 5;17 | no | no | P0A7J7 |
| 48 | TIKITQTR | Ac-WGGGGRTIKITQTR | 5;18 | 4;13 | no | P0AG51 |
| 49 | VAKSNVPALEACPQK | Ac-WGGGGRVAKSNVPALEACPQK(N3) | 5;19 | no | no | P0A7S3 |
| 50 | SNVPALEACPQKR | Ac-WGGGGRSNVPALEACPQKR | 5;20 | 4;14 | 6;12 | P0A7S3 |
| 51 | SKYGVK | Ac-WGGGGRSKYGVK(N3) | 6;11 | no | no | P0A7S3 |
| 52 | DEADEKDAIATVNK | Ac-WGGGGRDEADEKDAIATVNK(N3) | 6;12 | no | no | P0AG67 |
| 53 | VGFGYGKAR | Ac-WGGGGRVGFGYGKAR | 6;13 | 4;15 | no | P0A7W1 |
| 54 | KLMTEFNYNSVMQVPR | Ac-WGGGGRKLMTEFNYNSVMQVPR | 6;14 | 4;16 | no | P62399 |
| 55 | GLDITITTTAKSDEEGR | Ac-WGGGGRGLDITITTTAKSDEEGR | 6;15 | 5;9 | no | P62399 |
| 56 | QGYPIGCKVTLR | Ac-WGGGGRQGYPIGCKVTLR | 6;16 | 5;10 | no | P62399 |
| 57 | GLSAKSFDGR | Ac-WGGGGRGLSAKSFDGR | 6;17 | 5;11 | no | P62399 |
| 58 | SVAGFKIR | Ac-WGGGGRSVAGFKIR | 6;18 | 5;12 | no | P62399 |
| 59 | MYAVFQSGGKQHR | Ac-WGGGGRMYAVFQSGGKQHR | 6;19 | 5;13 | no | P0AG48 |
| 60 | KQQGHR | Ac-WGGGGRKQQGHR | 6;20 | 5;14 | no | P0AG48 |
| 61 | QLGEDPWVAIAKR | Ac-WGGGGRQLGEDPWVAIAKR | 7;11 | 5;15 | no | P0AG67 |
| 62 | YPEGTKLTGR | Ac-WGGGGRYPEGTKLTGR | 7;12 | 5;16 | no | P0AG67 |
| 63 | FWVESEKR | Ac-WGGGGRFWVESEKR | 7;13 | 6;9 | no | P0A7M2 |
| 64 | SHALNATKR | Ac-WGGGGRSHALNATKR | 4;7;14;15 | 6;10 | no | P0A7M2 |
| 65 | VSAKGMR | Ac-WGGGGRVSAKGMR | 7;15 | 6;11 | no | P0A7M2 |
| 66 | KDIHPK | Ac-WGGGGRKDIHPK(N3) | 7;16 | no | no | P0A7M9 |
| 67 | SHPFYTGKLR | Ac-WGGGGRSHPFYTGKLR | 7;17 | 6;12 | no | P0A7N1 |
| 68 | IGSTIKTDR | Ac-WGGGGRIGSTIKTDR | 7;18 | 6;13 | no | P0A7N1 |
| 69 | NFGKHPVTPWGVQTK | Ac-WGGGGRNFGKHPVTPWGVQTK(N3) | 7;19 | no | no | P60422 |
| 70 | HPVTPWGVQTKGK | Ac-WGGGGRHPVTPWGVQTKGK(N3) | 7;20 | no | no | P60422 |
| 71 | HVVKVVNPELHK | Ac-WGGGGRHVVKVVNPELHK(N3) | 8;11 | no | no | P60422 |
| 72 | HIGGGHKQAYR | Ac-WGGGGRHIGGGHKQAYR | 8;12 | 6;14 | no | P60422 |
| 73 | YILAPKGLK | Ac-WGGGGRYILAPKGLK(N3) | 8;13 | no | no | P60422 |
| 74 | DLETQSQDGTFDKLTK | Ac-WGGGGRDLETQSQDGTFDKLTK(N3) | 8;14 | no | no | P0A7V0 |
| 75 | DMLKAGVHFGHQTR | Ac-WGGGGRDMLKAGVHFGHQTR | 8;15 | 6;15 | no | P0A7V0 |
| 76 | MTDKLTSLR | Ac-WGGGGRMTDKLTSLRGGGG-Am | 8;16 | 6;16 | no | P0A870 |
| 77 | QPHAKGR | Ac-WGGGGRQPHAKGRGGGG-Am | 8;17 | 7;9 | no | P25888 |
| 78 | DYSKYLNIR | Ac-WGGGGRDYSKYLNIRGGGG-Am | 8;18 | 7;10 | no | P25888 |
| 79 | ASFDKANR | Ac-WGGGGRASFDKANRGGGG-Am | 8;19 | 7;11 | no | P0A715 |
| 80 | EGGNEKVILCDR | Ac-WGGGGREGGNEKVILCDRGGGG-Am | 8;20 | 7;12 | no | P0A715 |
| 81 | MTGRELKPHDR | Ac-WGGGGRMTGRELKPHDRGGGG-Am | 9;11 | 7;13 | no | P0A9Q1 |
| 82 | VVNSKEDIR | Ac-WGGGGRVVNSKEDIRGGGG-Am | 9;12 | 7;14 | no | P0A836 |
| 83 | EAEKYANPIPSR | Ac-WGGGGREAEKYANPIPSRGGGG-Am | 9;13 | 7;15 | no | P21499 |
| 84 | SMKQAIYDPENR | Ac-WGGGGRSMKQAIYDPENRGGGG-Am | 9;14 | 7;16 | no | P21499 |
| 85 | VGAATEVEMKEK | Ac-WGGGGRVGAATEVEMKEK(N3) | 9;15 | no | no | P0A6F5 |
| 86 | LWDKETLEK | Ac-WGGGGRLWDKETLEK(N3) | 9;16 | no | no | P0A7D7 |
| 87 | LSYDTEASIAKAK | Ac-WGGGGRLSYDTEASIAKAK(N3) | 9;17 | no | no | P0A870 |
| 88 | QMKAIDDLVK | Ac-WGGGGRQMKAIDDLVK(N3) | 9;18 | no | no | P0A715 |
| 89 | ESVLPKAVTAR | Ac-WGGGGRESVLPKAVTAR | 9;19 | 8;9 | no | P27302 |
| 90 | EKLQER | Ac-WGGGGREKLQER | 9;20 | 8;10 | no | P0A6F5 |
| 91 | AQSLKEIK | Ac-WGGGGRAQSLKEIK(N3) | 10;11 | no | no | P00956 |
| 92 | GEVKGK | Ac-WGGGGRGEVKGK(N3) | 10;12 | no | no | P0ABB0 |
| 93 | QLDHGQKVTELLK | Ac-WGGGGRQLDHGQKVTELLK(N3) | 10;13 | no | no | P0ABB0 |
| 94 | GLKVALSK | Ac-WGGGGRGLKVALSK(N3) | 10;14 | no | no | P00956 |
| 95 | DVKFGNDAR | Ac-WGGGGRDVKFGNDAR | 10;15 | 8;11 | no | P0A6F5 |
| 96 | DAAAAVGKAVAER | GGGG-AmRDAAAAVGKAVAER | 10;16 | 8;12 | no | P0C018 |
| 97 | KLQLVGVGYR | GGGG-AmRKLQLVGVGYR | 10;17 | 8;13 | no | P0AG55 |
| 98 | VAVIKAVR | GGGG-AmRVAVIKAVR | 10;18 | 8;14 | no | P0A7K2 |
| 99 | MQKQAELYR | Ac-WGGGGRMQKQAELYR | 10;19 | 8;15 | no | P0A7D7 |
| 100 | ATIDGLENMNSPEMVAAKR | Ac-WGGGGRATIDGLENMNSPEMVAAKR | 10;20 | 8;16 | no | P0A7W1 |
| 101 | QALELPR | Ac-WGGGGRQALELPRGGGG-Am | no | no | 1;7 | P0A850 |
| 102 | SQAIEGLVK | Ac-WGGGGRSQAIEGLVKGGGG-Am | no | no | 1;8 | P0A850 |
| 103 | SELVNVAK | Ac-WGGGGRSELVNVAKGGGG-Am | no | no | 1;10 | P0A850 |
| 104 | MTEAMK | Ac-WGGGGRMTEAMKGGGG-Am | no | no | 1;11 | P37095 |
| 105 | IDGLGIK | Ac-WGGGGRIDGLGIKGGGG-Am | no | no | 1;12 | P37095 |
| 106 | LPLAEFHR | Ac-WGGGGRLPLAEFHRGGGG-Am | no | no | 1;13 | P37095 |
| 107 | QTAFMDSMK | Ac-WGGGGRQTAFMDSMKGGGG-Am | no | no | 2;7 | P37095 |
| 108 | ITLSTQPADAR | Ac-WGGGGRITLSTQPADARGGGG-Am | no | no | 2;8 | P37095 |
| 109 | GITFDSGGYSIK | Ac-WGGGGRGITFDSGGYSIKGGGG-Am | no | no | 2;9 | P37095 |
| 110 | EQGYMGLHTVGR | Ac-WGGGGREQGYMGLHTVGRGGGG-Am | no | no | 2;10 | P37095 |
| 111 | EAGQAK | Ac-WGGGGREAGQAKGGGG-Am | no | no | 2;11 | P27302 |
| 112 | LTAEGVK | Ac-WGGGGRLTAEGVKGGGG-Am | no | no | 2;12 | P27302 |
| 113 | QDAAYR | Ac-WGGGGRQDAAYRGGGG-Am | no | no | 2;13 | P27302 |
| 114 | ALSMDAVQK | Ac-WGGGGRALSMDAVQKGGGG-Am | no | no | 3;7 | P27302 |
| 115 | QNLAQQER | Ac-WGGGGRQNLAQQERGGGG-Am | no | no | 3;8 | P27302 |
| 116 | QDGPTALILSR | Ac-WGGGGRQDGPTALILSRGGGG-Am | no | no | 3;9 | P27302 |
| 117 | AVTDKPSLLMCK | Ac-WGGGGRAVTDKPSLLMCKGGGG-Am | no | no | 3;10 | P27302 |
| 118 | VLDAAVAGK | Ac-WGGGGRVLDAAVAGKGGGG-Am | no | no | 3;11 | P0A6P1 |
| 119 | TGAGMMDCK | Ac-WGGGGRTGAGMMDCKGGGG-Am | no | no | 3;12 | P0A6P1 |
| 120 | AGNVAADGVIK | Ac-WGGGGRAGNVAADGVIKGGGG-Am | no | no | 3;13 | P0A6P1 |
| 121 | MAEITASLVK | Ac-WGGGGRMAEITASLVKGGGG-Am | no | no | 4;7 | P0A6P1 |
| 122 | EAAPAK | Ac-WGGGGREAAPAKGGGG-Am | no | no | 4;8 | P0A9Q7 |
| 123 | AELGIPK | Ac-WGGGGRAELGIPKGGGG-Am | no | no | 4;9 | P0A9Q7 |
| 124 | ALIVTDR | Ac-WGGGGRALIVTDRGGGG-Am | no | no | 4;10 | P0A9Q7 |
| 125 | AVQDVILK | Ac-WGGGGRAVQDVILKGGGG-Am | no | no | 4;11 | P0A9Q7 |
| 126 | AAALAAADAR | Ac-WGGGGRAAALAAADARGGGG-Am | no | no | 4;12 | P0A9Q7 |
| 127 | YPLISELK | Ac-WGGGGRYPLISELKGGGG-Am | no | no | 4;13 | P0A9Q7 |
| 128 | YNANDNPTK | Ac-WGGGGRYNANDNPTKGGGG-Am | no | no | 5;7 | P0A9Q7 |
| 129 | TGDTISGK | Ac-WGGGGRTGDTISGKGGGG-Am | no | no | 5;8 | P0AG30 |
| 130 | MNLTELK | Ac-WGGGGRMNLTELKGGGG-Am | no | no | 5;9 | P0AG30 |
| 131 | VFPAIDYNR | Ac-WGGGGRVFPAIDYNRGGGG-Am | no | no | 5;10 | P0AG30 |
| 132 | GTGNMELHLSR | Ac-WGGGGRGTGNMELHLSRGGGG-Am | no | no | 5;11 | P0AG30 |
| 133 | AGNDANR | Ac-WGGGGRAGNDANRGGGG-Am | no | no | 5;12 | P00864 |
| 134 | VLGETIK | Ac-WGGGGRVLGETIKGGGG-Am | no | no | 5;13 | P00864 |
| 135 | ATDLFLK | Ac-WGGGGRATDLFLKGGGG-Am | no | no | 6;7 | P00864 |
| 136 | MVEVNACLK | Ac-WGGGGRMVEVNACLKGGGG-Am | no | no | 6;8 | P00864 |
| 137 | LPVEFVPVR | Ac-WGGGGRLPVEFVPVRGGGG-Am | no | no | 6;9 | P00864 |
| 138 | MNEQYSALR | Ac-WGGGGRMNEQYSALRGGGG-Am | no | no | 6;10 | P00864 |
| 139 | SLTEIK | Ac-WGGGGRSLTEIKGGGG-Am | no | no | 6;11 | P0A7Z4 |
| 140 | HWDQQK | Ac-WGGGGRHWDQQKGGGG-Am | no | no | 6;13 | P25665 |
| 141 | AQESYWAGNSTR | Ac-WGGGGRAQESYWAGNSTRGGGG-Am | no | no | 7; | P25665 |

| Supplementary Table 2; # of unique XL sites | | | total number | | | | | correct number | | | | | false number | | | | | FDR | | | | |
|---|---|---|---|---|---|---|---|---|---|---|---|---|---|---|---|---|---|---|---|---|---|---|
| | | | MeroX | Annika | XlinkX | pLink | MaxLynx | MeroX | Annika | XlinkX | pLink | MaxLynx | MeroX | Annika | XlinkX | pLink | MaxLynx | MeroX | Annika | XlinkX | pLink | MaxLynx |
| main library | DSSO | replicate 1 | 638 | 581 | 463 | 420 | 585 | 593 | 563 | 447 | 400 | 568 | 45 | 18 | 16 | 20 | 17 | 7.1% | 3.1% | 3.5% | 4.8% | 2.9% |
| | | replicate 2 | 612 | 599 | 510 | 385 | 608 | 580 | 583 | 482 | 373 | 600 | 32 | 16 | 28 | 12 | 8 | 5.2% | 2.7% | 5.5% | 3.1% | 1.3% |
| | | replicate 3 | 724 | 715 | 605 | 564 | 711 | 688 | 697 | 579 | 541 | 695 | 36 | 18 | 26 | 23 | 16 | 5.0% | 2.5% | 4.3% | 4.1% | 2.3% |
| | | average | 658 | 632 | 526 | 456 | 635 | 620 | 614 | 503 | 438 | 621 | 38 | 17 | 23 | 18 | 14 | 5.7% | 2.7% | 4.4% | 4.0% | 2.2% |
| | | standard deviation | 59 | 73 | 72 | 95 | 67 | 59 | 72 | 68 | 90 | 66 | 7 | 1 | 6 | 6 | 5 | - | - | - | - | - |
| | DSBU | replicate 1 | 804 | 594 | 590 | 412 | 764 | 752 | 572 | 528 | 398 | 741 | 52 | 22 | 62 | 14 | 23 | 6.5% | 3.7% | 10.5% | 3.4% | 3.0% |
| | | replicate 2 | 746 | 561 | 550 | 428 | 676 | 697 | 547 | 512 | 412 | 666 | 49 | 14 | 38 | 16 | 10 | 6.6% | 2.5% | 6.9% | 3.7% | 1.5% |
| | | replicate 3 | 751 | 520 | 546 | 455 | 682 | 695 | 508 | 512 | 434 | 667 | 56 | 12 | 34 | 21 | 15 | 7.5% | 2.3% | 6.2% | 4.6% | 2.2% |
| | | average | 767 | 558 | 562 | 432 | 707 | 715 | 542 | 517 | 415 | 691 | 52 | 16 | 45 | 17 | 16 | 6.8% | 2.9% | 7.9% | 3.9% | 2.3% |
| | | standard deviation | 32 | 37 | 24 | 22 | 49 | 32 | 32 | 9 | 18 | 43 | 4 | 5 | 15 | 4 | 7 | - | - | - | - | - |
| | CDI | replicate 1 | 86 | 68 | 44 | 39 | 83 | 84 | 62 | 40 | 37 | 79 | 2 | 6 | 4 | 2 | 4 | 2.3% | 8.8% | 9.1% | 5.1% | 4.8% |
| | | replicate 2 | 83 | 65 | 44 | 47 | 61 | 77 | 63 | 40 | 45 | 58 | 6 | 2 | 4 | 2 | 3 | 7.2% | 3.1% | 9.1% | 4.3% | 4.9% |
| | | replicate 3 | 73 | 54 | 37 | 43 | 54 | 67 | 47 | 35 | 39 | 50 | 6 | 7 | 2 | 4 | 4 | 8.2% | 13.0% | 5.4% | 9.3% | 7.4% |
| | | average | 81 | 62 | 42 | 43 | 66 | 76 | 57 | 38 | 40 | 62 | 5 | 5 | 3 | 3 | 4 | 5.9% | 8.0% | 8.0% | 6.2% | 5.6% |
| | | standard deviation | 7 | 7 | 4 | 4 | 15 | 9 | 9 | 3 | 4 | 15 | 2 | 3 | 1 | 1 | 1 | - | - | - | - | - |
| enrichable sublibrary | DSSO | replicate 1 | 426 | 400 | 392 | 393 | 469 | 391 | 383 | 367 | 362 | 454 | 35 | 17 | 25 | 31 | 15 | 8.2% | 4.3% | 6.4% | 7.9% | 3.2% |
| | | replicate 2 | 435 | 423 | 384 | 405 | 478 | 398 | 399 | 358 | 370 | 461 | 37 | 24 | 26 | 35 | 17 | 8.5% | 5.7% | 6.8% | 8.6% | 3.6% |
| | | replicate 3 | 392 | 352 | 323 | 293 | 423 | 354 | 338 | 302 | 276 | 411 | 38 | 14 | 21 | 17 | 12 | 9.7% | 4.0% | 6.5% | 5.8% | 2.8% |
| | | average | 418 | 392 | 366 | 364 | 457 | 381 | 373 | 342 | 336 | 442 | 37 | 18 | 24 | 28 | 15 | 8.8% | 4.7% | 6.6% | 7.6% | 3.2% |
| | | standard deviation | 23 | 36 | 38 | 61 | 30 | 24 | 32 | 35 | 52 | 27 | 2 | 5 | 3 | 9 | 3 | - | - | - | - | - |
| | DSBSO | replicate 1 | 409 | 370 | 363 | 387 | 416 | 375 | 362 | 334 | 350 | 408 | 34 | 8 | 29 | 37 | 8 | 8.3% | 2.2% | 8.0% | 9.6% | 1.9% |
| | | replicate 2 | 407 | 361 | 347 | 382 | 407 | 382 | 355 | 323 | 349 | 398 | 25 | 6 | 24 | 33 | 9 | 6.1% | 1.7% | 6.9% | 8.6% | 2.2% |
| | | replicate 3 | 378 | 347 | 299 | 291 | 382 | 361 | 334 | 276 | 271 | 376 | 17 | 13 | 23 | 20 | 6 | 4.5% | 3.7% | 7.7% | 6.9% | 1.6% |
| | | average | 398 | 359 | 336 | 353 | 402 | 373 | 350 | 311 | 323 | 394 | 25 | 9 | 25 | 30 | 8 | 6.4% | 2.5% | 7.5% | 8.5% | 1.9% |
| | | standard deviation | 17 | 12 | 33 | 54 | 18 | 11 | 15 | 31 | 45 | 16 | 9 | 4 | 3 | 9 | 2 | - | - | - | - | - |
| acidic library | ADH, XL in separate gorups | replicate 1 | 26 | - | 22 | 111 | 95 | 21 | - | 21 | 104 | 90 | 5 | - | 1 | 7 | 5 | 19.2% | - | 4.5% | 6.3% | 5.3% |
| | | replicate 2 | 36 | - | 18 | 88 | 95 | 29 | - | 17 | 82 | 86 | 7 | - | 1 | 6 | 9 | 19.4% | - | 5.6% | 6.8% | 9.5% |
| | | replicate 3 | 32 | - | 53 | 102 | 83 | 26 | - | 44 | 91 | 78 | 6 | - | 9 | 11 | 5 | 18.8% | - | 17.0% | 10.8% | 6.0% |
| | | average | 31 | - | 31 | 100 | 91 | 25 | - | 27 | 92 | 85 | 6 | - | 4 | 8 | 6 | 19.1% | - | 11.8% | 8.0% | 7.0% |
| | | standard deviation | 5 | - | 19 | 12 | 7 | 4 | - | 15 | 11 | 6 | 1 | - | 5 | 3 | 2 | - | - | - | - | - |
| | ADH, XL in pooled group | replicate 1 | 46 | - | 85 | 220 | - | 32 | - | 81 | 217 | - | 14 | - | 4 | 3 | - | 30.4% | - | 4.7% | 1.4% | - |
| | | replicate 2 | 37 | - | 2 | 226 | - | 31 | - | 0 | 224 | - | 6 | - | 2 | 2 | - | 16.2% | - | 100.0% | 0.9% | - |
| | | replicate 3 | 44 | - | 3 | 209 | - | 34 | - | 1 | 202 | - | 10 | - | 2 | 7 | - | 22.7% | - | 66.7% | 3.3% | - |
| | | average | 42 | - | 30 | 218 | - | 32 | - | 27 | 214 | - | 10 | - | 3 | 4 | - | 23.6% | - | 8.9% | 1.8% | - |
| | | standard deviation | 5 | - | 48 | 9 | - | 2 | - | 46 | 11 | - | 4 | - | 1 | 3 | - | - | - | - | - | - |
| | DHSO, XL in separate gorups | replicate 1 | 76 | 94 | 89 | 68 | 93 | 74 | 90 | 74 | 59 | 88 | 2 | 4 | 15 | 9 | 5 | 2.6% | 4.3% | 16.9% | 13.2% | 5.4% |
| | | replicate 2 | 92 | 65 | 79 | 68 | 104 | 86 | 62 | 66 | 63 | 96 | 6 | 3 | 13 | 5 | 8 | 6.5% | 4.6% | 16.5% | 7.4% | 7.7% |
| | | replicate 3 | 71 | 65 | 65 | 76 | 99 | 67 | 61 | 57 | 61 | 84 | 4 | 4 | 8 | 15 | 15 | 5.6% | 6.2% | 12.3% | 19.7% | 15.2% |
| | | average | 80 | 75 | 78 | 71 | 99 | 76 | 71 | 66 | 61 | 89 | 4 | 4 | 12 | 10 | 9 | 5.0% | 4.9% | 15.5% | 13.7% | 9.5% |
| | | standard deviation | 11 | 17 | 12 | 5 | 6 | 10 | 16 | 9 | 2 | 6 | 2 | 1 | 4 | 5 | 5 | - | - | - | - | - |
| | DHSO, XL in pooled group | replicate 1 | 141 | 180 | 179 | 84 | - | 132 | 178 | 171 | 83 | - | 9 | 2 | 8 | 1 | - | 6.4% | 1.1% | 4.5% | 1.2% | - |
| | | replicate 2 | 132 | 204 | 152 | 145 | - | 126 | 195 | 144 | 142 | - | 6 | 9 | 8 | 3 | - | 4.5% | 4.4% | 5.3% | 2.1% | - |
| | | replicate 3 | 93 | 176 | 157 | 161 | - | 90 | 173 | 146 | 154 | - | 3 | 3 | 11 | 7 | - | 3.2% | 1.7% | 7.0% | 4.3% | - |
| | | average | 122 | 187 | 163 | 130 | - | 116 | 182 | 154 | 126 | - | 6 | 5 | 9 | 4 | - | 4.9% | 2.5% | 5.5% | 2.8% | - |
| | | standard deviation | 26 | 15 | 14 | 41 | - | 23 | 12 | 15 | 38 | - | 3 | 4 | 2 | 3 | - | - | - | - | - | - |

| **Supplementary Table 3**: XL search engine settings, if not given different for specific results | **MeroX** | **MS Annika** | **XlinkX** | **pLink** | **MaxLynx** |
|---|---|---|---|---|---|
| software suite/version | standalone/ v 2.0.1.4 | in PD 2.5, v 1.2.17259 | part of PD 2.5 | standalone/ v2.3.9 | part of MaxQuant v 2.0.2.0 |
| search mode | RISEUP (Quadratic mode for ADH) | combined | - | conventional XL | - |
| MS1 tolerance | 5 ppm | 5 ppm | 5 ppm | 5 ppm | 5 ppm |
| MS2 tolerance | 10 ppm | 10 ppm | 10 ppm | 10 ppm | 10 ppm |
| enzymatic cleavage | K, R blocked by P | | | | |
| missed cleavages | 3 | 3 | 3 | 3 | 3 |
| crosslink modification (DSSO, DSBU,CDI, DSBSO) | K to K | K to K | K to K | K to K | K to K |
| crosslink modification (DHSO, ADH) | D/E to D/E | D/E to D/E | D/E to D/E | D/E to D/E | D/E to D/E |
| PTMs | Carbamidomethyl (C, Static), Oxidation (M, Dynamic) | | | | |
| minimal peptide length | 6 | 6 | 6 | 6 | 6 |
| minimal score | -1 | 0 | 0/ delta score 4 | - | 0 / min partial score 10 |
| FDR | 1% | | | | |
| separate inter- intra-link FDR | active | inactive | inactive | active | active |
| database | custom E.coli ribosome shotgun, 171 sequences | | | | |
